# Quantitative drug susceptibility testing for M. tuberculosis using unassembled sequencing data and machine learning

**DOI:** 10.1101/2021.09.14.458035

**Authors:** The CRyPTIC consortium, Alexander S Lachapelle

**Affiliations:** University of Oxford

## Abstract

There remains a clinical need for better approaches to rapid drug susceptibility testing in view of the increasing burden of multidrug resistant tuberculosis. Binary susceptibility phenotypes only capture changes in minimum inhibitory concentration when these cross the critical concentration, even though other changes may be clinically relevant. We developed a machine learning system to predict minimum inhibitory concentration from unassembled whole-genome sequencing data for 13 anti-tuberculosis drugs. We trained, validated and tested the system on 10,859 isolates from the CRyPTIC dataset. Essential agreement rates (predicted MIC within one doubling dilution of observed MIC) were above 92% for first-line drugs, 91% for fluoroquinolones and aminoglycosides, and 90% for new and repurposed drugs, albeit with a significant drop in performance for the very few phenotypically resistant isolates in the latter group. To further validate the model in the absence of external MIC datasets, we predicted MIC and converted values to binary for an external set of 15,239 isolates with binary phenotypes, and compare their performance against a previously validated mutation catalogue, the expected performance of existing molecular assays, and World Health Organization Target Product Profiles. The sensitivity of the model on the external dataset was greater than 90% for all drugs except ethionamide, clofazimine and linezolid. Specificity was greater than 95% for all drugs except ethambutol, ethionamide, bedaquiline, delamanid and clofazimine. The proposed system can provide quantitative susceptibility phenotyping to help guide antimicrobial therapy, although further data collection and validation are required before machine learning can be used clinically for all drugs.

## Background

In 2020, 10 million individuals fell ill with tuberculosis and 1.5 million died from the infection^1^. The problem of multidrug-resistant tuberculosis (MDR-TB) - defined as resistance to isoniazid and rifampicin - has been described by the World Health Organization (WHO) as a global health crisis.^1^ Despite advances in diagnostics and treatment, MDR-TB remains under-detected and treatment success remains stubbornly below 60% globally.^1,2^ The SARS-CoV-2 pandemic has likely set back the progress that has been made by years.^3^

The WHO has called for universal tuberculosis drug susceptibility testing (DST),^4^ but two major barriers remain for universal DST in the face of rapidly evolving resistance. First, culture-based DST is too slow, expensive, and technically challenging to offer a realistic solution. Novel molecular assays have been developed to rapidly detect resistance to a subset of first- and second-line drugs, but are limited in the number of resistance-conferring mutations they can detect,^5^ constraining their sensitivity. Some countries now rely on whole-genome sequencing (WGS) to identify susceptibility to first-line drugs,^6^ but suffer the same limitations in the number of resistance-conferring mutations detected.. Second, DST is largely based on binary phenotypes, where susceptible isolates are differentiated from resistant ones based on critical concentrations of antibiotic that are not always firmly relatable to clinical outcome. Furthermore, sub-critical concentration elevations in minimum inhibitory concentration (MIC) that are of potential clinical significance cannot be identified.

Machine learning algorithms have accurately predicted binary susceptibility from whole genome sequencing data in *M. tuberculosis*^7–10^, and predicted MICs in other pathogens like *Salmonella*^11^, *N. gonorrhea*^12^, *S. aureus*^13^, and *E. coli*^14^. The difficulty in generating MIC data for M. tuberculosis has meant that machine learning approaches have yet to be applied to the problem of predicting MICs from WGS data for this major pathogen.

Here, we assess how well a machine learning system can predict MIC for 13 anti-tuberculosis drugs, including new and repurposed drugs like bedaquiline and delamanid, using genome-wide genomic features. We leverage a new, recently published dataset from our CRyPTIC consortium, and adapt previously described extreme gradient boosting machine models for use on genome-wide *k*-mers directly from unassembled sequencing reads. We then assess whether MIC predictions can be converted to predict susceptibility or resistance for isolates from an independent dataset with binary Mycobacterial Growth Indicator Tubes (MGIT) based results.

## Results

### Dataset characteristics

The CRyPTIC dataset included MIC phenotypes for 10,859 isolates from 22 countries. Lineages 1 to 4, 6 and *Mycobacterium bovis* were represented, with lineage 4 (50%, 5,436/10,859) and lineage 2 (35%, 3,745/10,859) the most common. 28% of samples (3,033/10,859) were MDR. MIC phenotypes were collected using a 96-well broth microdilution plate, and were available for three first-line antibiotics (isoniazid, rifampicin, ethambutol), plus rifabutin, two fluoroquinolones (moxifloxacin, levofloxacin), two injectable agents (amikacin, kanamycin), ethionamide, and four new or repurposed drugs (bedaquiline, linezolid, clofazimine, delamanid). Pyrazinamide was not present on the microdilution plate for technical reasons. We present results for kanamycin and rifabutin exclusively in the supplementary appendix as they are less commonly prescribed.

### Minimum inhibitory concentration prediction

We developed a *k-*mer-based, hypothesis-free, genome-wide supervised machine learning algorithm to predict MIC for 13 antibiotics (Methods). To assess the performance of the machine learning system on the widest possible set of antibiotics, it was initially trained on 75% of the CRyPTIC dataset (8,146 randomly selected isolates) (Figure S1). Predictions were made for the remaining 25% (2,713 isolates). We generated confidence matrices for each drug (Figure 2) and calculated essential agreement rates, or when the MIC is correctly predicted within one doubling dilution (Table 2).

**Figure 1:**
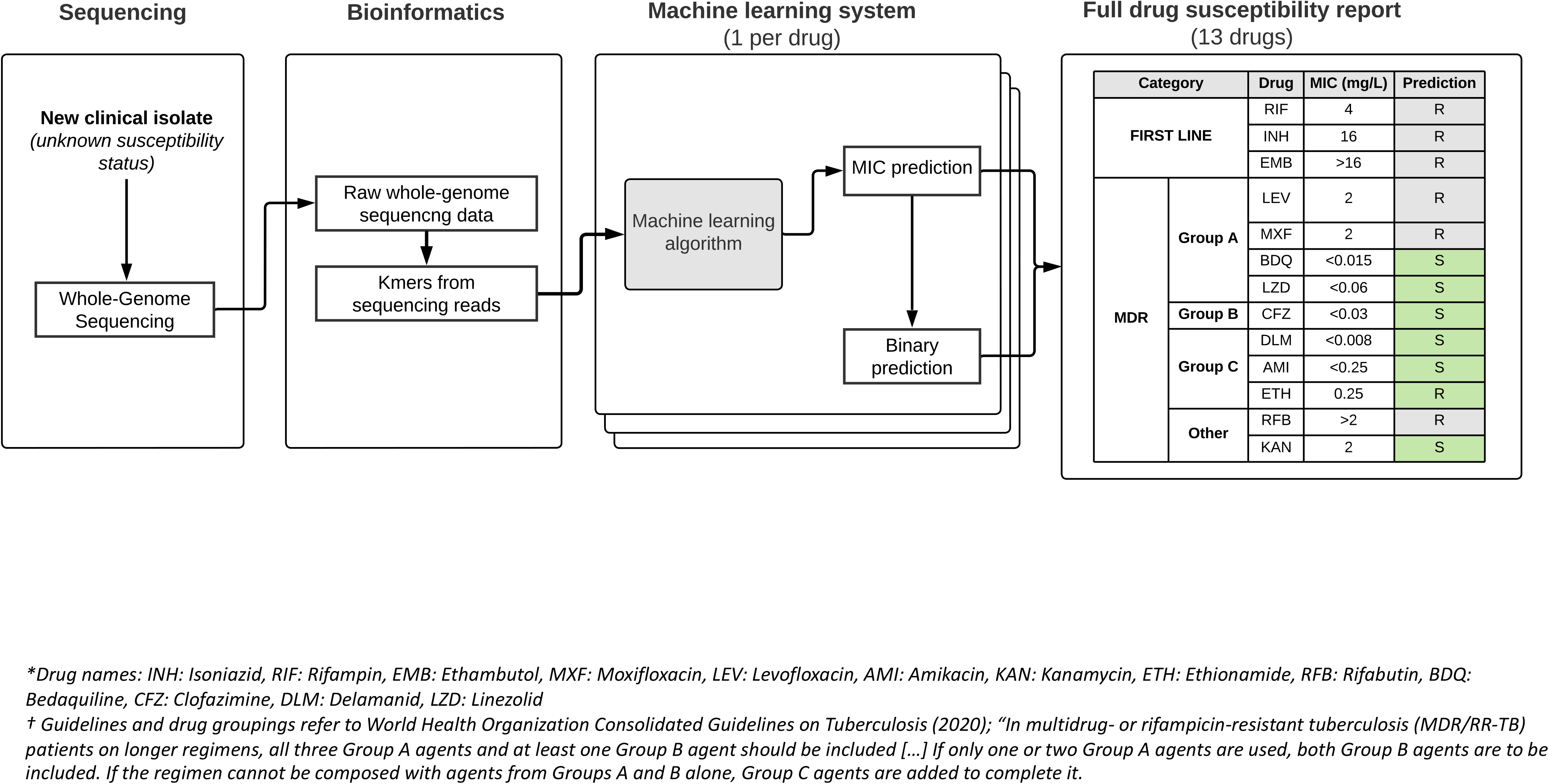
Overview of the machine learning prediction system and prediction workflow

**Figure 2:**
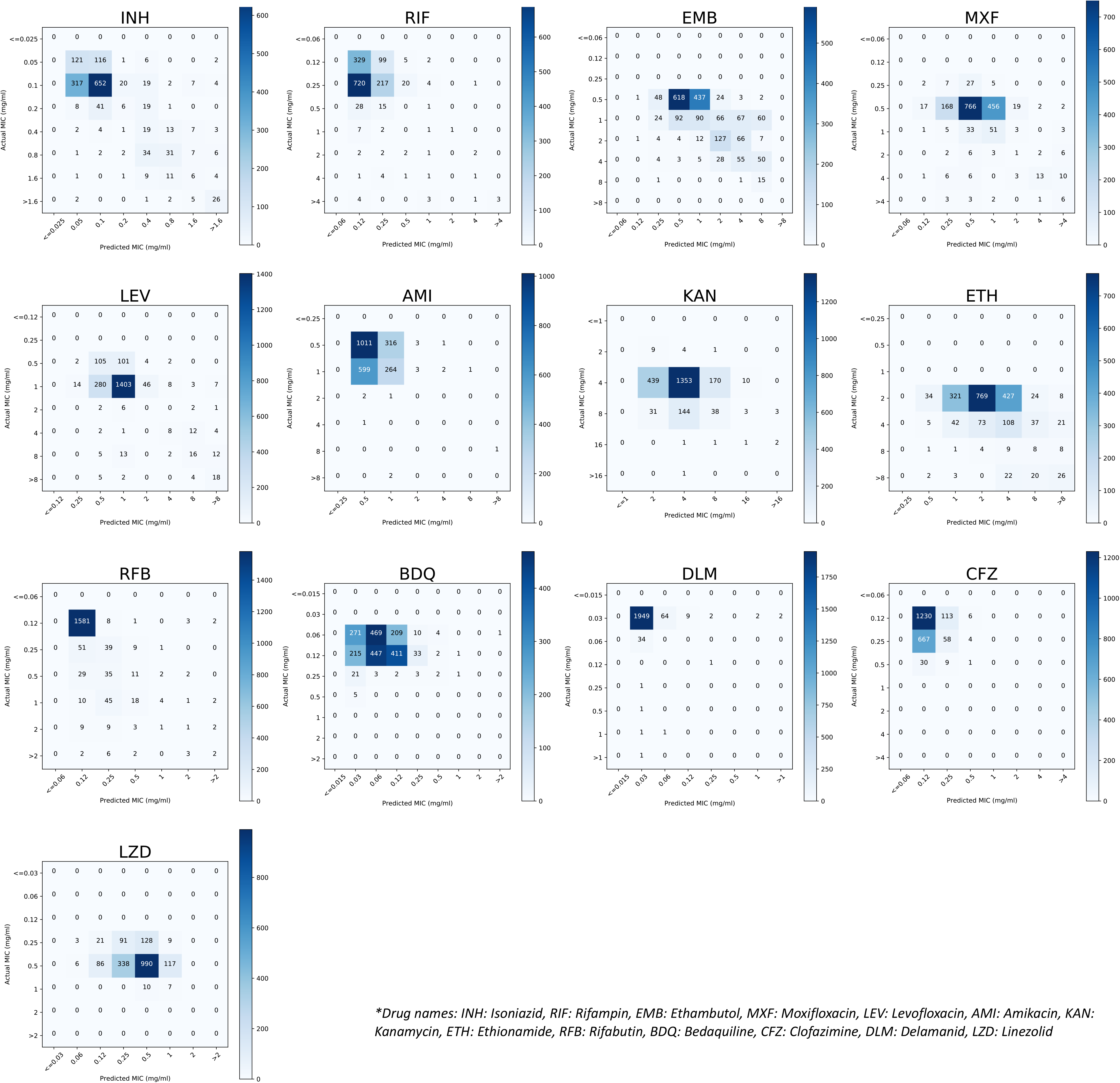
Predicted and actual mimimum inhibitory concentrations for 13 antituberculosis drugs in the CRyPTIC dataset

**Figure 3:**
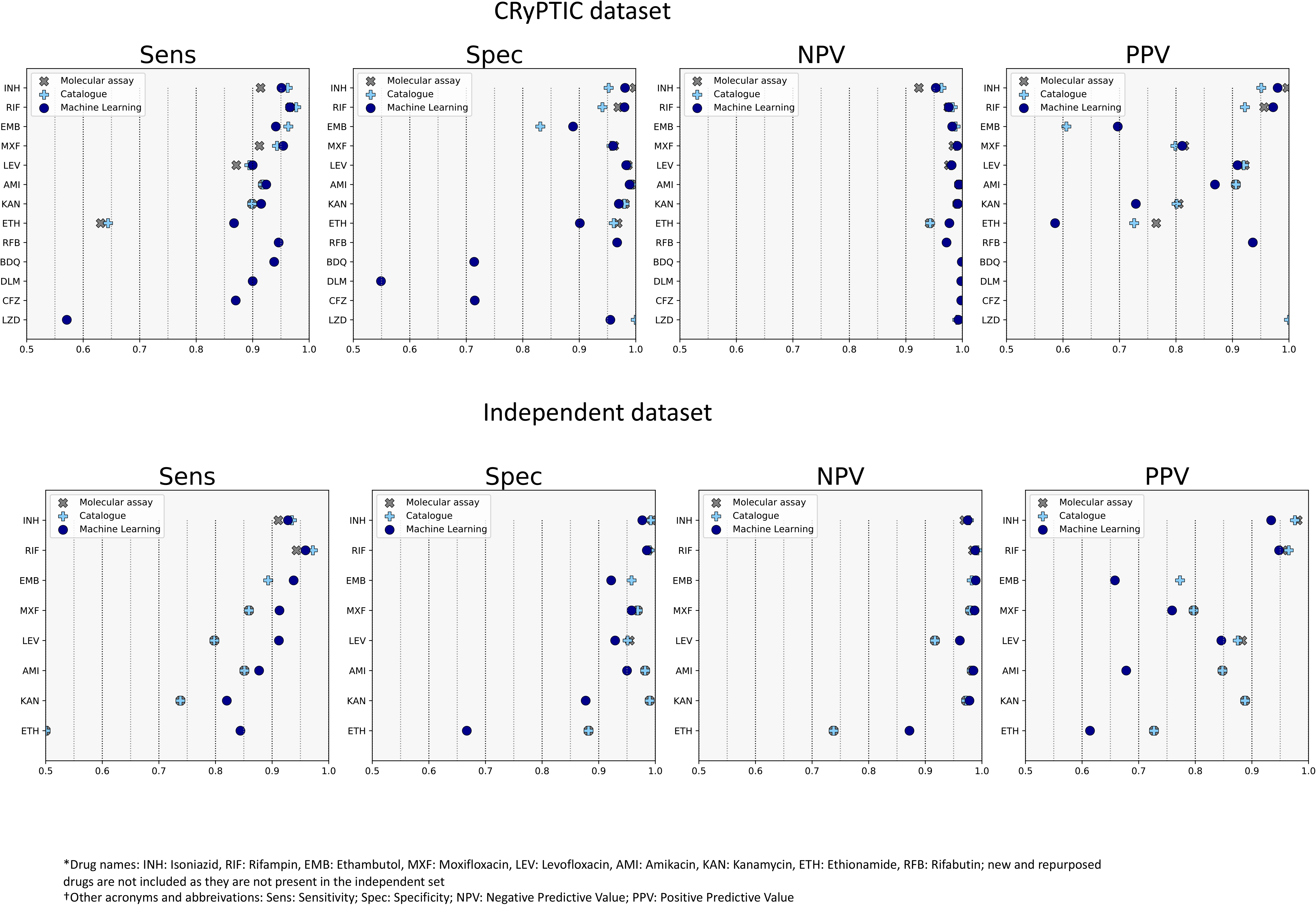
Performance of the machine learning prediction on the CRyPTIC (a) and independent (b) test sets after MIC predictions were converted to binary predictions, and comparison to the catalogue and molecular assay

For the first-line drugs, the essential agreement (EA) rate was 94.2% for isoniazid (2,242/2,380), 95.0% for ethambutol (1,811/1,907), and 92.9% for rifampicin (2,086/2,246) (Table 2). We compared EA for phenotypically resistant (MIC above epidemiological cutoff^15^) and phenotypically susceptible (MIC below epidemiological cutoff). There was no statistically significant difference in EA for ethambutol and rifampicin. There was a 7.3% decrease in EA for phenotypically resistant samples compared to susceptible isolates for isoniazid (97.8% *vs* 90.5%, p<0.001). Perfect agreement rates (exact MIC predicted correctly) were on average 30.1% lower than essential agreement rates across drugs (p<0.001) and are presented in the Supplementary Appendix (Table S2a).

EA was 94.9% for levofloxacin (1,837/1,936), 90.9% for moxifloxacin (1,544/1,699), 97.5% for amikacin (2,191/2,248), and 91.3% for ethionamide (2,017/2,210). EA was lower for phenotypically resistant than for phenotypically susceptible isolates for all drugs, with differences of 13.1%, 7.3%, 10.9%, and 20.0%, for levofloxacin, moxifloxacin, amikacin, and ethionamide, respectively (p<0.001). We note that there were fewer phenotypically resistant isolates for these four drugs: 310, 261, 158, and 309, respectively, in the CRyPTIC dataset - which could explain the challenge in lower accuracy. EA was above 93% for all new and repurposed drugs: 93.3% for bedaquiline (1,987/2,130), 96.1% for clofazimine (1,842/1,917), 93.6% for linezolid (1,689/1,805), and 96.4% (1,918/1,990). EA dropped significantly for phenotypically resistant isolates: 39.3% (11/28) for linezolid, 13.0% for clofazimine (3/23), 10.0% for delamanid (2/20), and 6.2% for bedaquiline (1/16), demonstrating the poor predictive performance for these drugs, with the high EA due to the majority of susceptible predictions. We observe the bimodal distribution of MICs for rifampicin, rifabutin, amikacin, and isoniazid, while MICs for ethambutol, kanamycin, moxifloxacin, and levofloxacin show a more unimodal distribution, correlating with less accurate predictions distributed across a broader range of the MIC confusion matrix (Figure 2) and consistent with previous findings in the CRyPTIC dataset^15^.

### Binarising M/Cs

After predicting MIC, we assessed whether the machine learning model could be used to predict binary susceptibility status, given binary predictions remain the standard by which most patients are treated. We begin by considering the MIC as a probability of resistance (i.e. increased MIC as increased probability of resistance), enabling us to calculate an area under the curve (AUC) metric (Table 3, Figure S2). For first-line drugs, AUC was 0.983 for isoniazid (95% CI 0.979-0.987), 0.992 for rifampicin (0.988-0.994), and 0.965 for ethambutol (0.957-0.973). For second-line drugs, AUC was 0.945 for levofloxacin (0.931-0.963), 0.970 for moxifloxacin (0.958-0.982), 0.970 for amikacin (0.953-0.984), 0.960 for kanamycin (0.941-0.975), and 0.939 for ethionamide (0.924-0.952). For new and repurposed drugs, AUCs were lower - 0.908 for bedaquiline (0.876-0.952), 0.879 for clofazimine (0.832-0.934), 0.781 for linezolid (0.695-0.866) and 0.785 for delamanid (0.711-0.861). However, we note that these four results are likely significant overestimations due to the very low number of true phenotypic positives in the test set (between 14 and 19). Receiver operating characteristic (ROC) curves for each drug are presented in the supplementary appendix (Figure S2).

We then converted the MIC predictions to binary predictions (Method). For first-line drugs, sensitivity was 95.0% for isoniazid, 99.2% for rifampicin, and 93.1% for ethambutol, with a specificity of 98.1%, 89.4%, and 97.3%, respectively. Sensitivity was 91.6% for levofloxacin, 95.0% for moxifloxacin, 91.8% for amikacin, and 87.4% for ethionamide, with a specificity of 96.1%, 96.1%, 99.1%, and 89.4%, respectively. For new and repurposed drugs, we find a sensitivity of 87.5% for bedaquiline, 82.6% for clofazimine, 57.1% for linezolid, and 75.0% for delamanid, with a specificity of 76.0%, 72.2%, 87.6%, and 72.4%, respectively (Table 3).

#### External validation and comparison to diagnostic standards

No dataset yet exists to validate the performance of MIC predictions for the full spectrum of drugs on the CRyPTIC plate. In order to measure the performance of our predictor on an external dataset, we assess how a model trained on the entire CRyPTIC dataset can predict binary resistance in a separate set of 15,577 isolates for first-line and second-line drugs from the Seq&Treat dataset. AUC was 0.971 for isoniazid (95%CI 0.967-0.975) and 0.981 for rifampicin (0.978-0.984), in both cases 1.1% lower than in CRyPTIC (p<0.01). Sensitivity was 93.6% for isoniazid and 96.0% for rifampicin, with a specificity of 99.2% and 99.0%, respectively. For second-line drugs, AUC was higher than CRyPTIC for levofloxacin (0.949), and lower for moxifloxacin (0.948), amikacin (0.924), and ethionamide (0.812). Sensitivity was 91.2%, 91.6%, 87.7%, and 90.6%, respectively, with a specificity of 92.5%, 95.1%, 94.9%, and 66.7%, respectively (Table 3).

We next compared predictions from the machine learning system with those from a validated mutation catalogue^16^ in the independent set. For first-line drugs, the catalogue had similar sensitivity (0.935 vs 0.936) with a higher specificity (0.992 vs 0.963) for isoniazid, and higher sensitivity (0.972 vs 0.960) and specificity (0.990 vs 0.984) for rifampicin. The machine learning system performed with much higher sensitivity for fluoroquinolones, with an increase in sensitivity for levofloxacin (91.2% vs 79.8%) and moxifoxacin (91.6% vs 85.9%), with slightly lower specificity (92.5% vs 95.1%, and 96.9 vs 95.1%, respectively). For amikacin, sensitivity was also higher (87.7% vs 85.1%), with lower specificity (94.9% vs 98.2%) (Table 3).

As most patients in the world have little or no access to phenotypic DST, we also compared the performance of the machine learning system against the expected combined performance of Xpert MTB/RIF and Xpert XDR for the independent set, in anticipation of its wider uptake to address the WHO’s call for universal DST. The sensitivity of the machine learning system was 24% higher than Xpert for isoniazid (93.6% vs 91.1%, p<0.001) and rifampicin (96.0% vs 94.3%, p<0.001), 6% higher for moxifloxacin (91.6% vs 85.9%, p<0.001), and 2% higher for amikacin (87.7% vs 85.1%, p=0.030). Specificity was no more than 1% lower for each drug, with the exception of isoniazid (96.3% vs 99.4%) and amikacin (94.9% vs 98.2%).

## Discussion

We assess for the first time the extent to which machine learning can predict minimum inhibitory concentration of tuberculosis isolates based exclusively on whole-genome sequencing data for 13 antituberculosis drugs, including new and repurposed drugs. We use extreme gradient boosting, previously demonstrated to perform well on other bacteria and for binary tuberculosis prediction, and a set of unassembled genome-wide *k-*mers from sequencing reads instead of the more common approach of using genomic mutations. We followed best practice guidance for studies evaluating the accuracy of rapid tuberculosis drug susceptibility testing (DST).^17^

We find that our machine learning model can predict MIC with an essential agreement rate greater than 93% for first-line drugs, and 90% for other drugs. The WHO target product profiles (TPP) for priority antituberculosis drugs, do not include thresholds and objectives for MIC prediction, focusing instead on binary metrics like sensitivity and specificity. However, essential agreements are similar to those reported for some other bacteria like *Salmonella*^11^, *N. gonorrhea*^12^, *S. aureus*^13^, and *E. coli*^14^.

MIC data lend themselves to the development for personalised dosage regimens in the future. While WHO-endorsed critical concentrations (CCs) have become widely used, elevations in MIC below or close to those CCs are not normally identified^18^, despite being potentially meaningful.

Current WHO guidelines for MDR-TB management recommend offering patients regimens including bedaquiline, linezolid, and potentially also clofazimine.^18^ Our sensitivity and specificity for these three drugs fall below WHO TPP requirements. Our results reflect the very low prevalence of resistance to these drugs, which also explains the high negative predictive values and low positive predictive values. As more resistant isolates are collected, the sensitivity and specificity of the machine learning system will almost certainly increase, and negative predictive value decrease a little, as seen for other drugs. Nevertheless, a test with a high negative predictive value, lower positive predictive value, and sensitivity of 70% would still be helpful to clinicians.^19^ Even imperfect test performance could still play a key role in preventing the amplification and dissemination of resistance.

For drugs where the WHO-endorsed molecular GeneXpert assay is available (rifampicin for Xpert MTB/RIF, and isoniazid, fluoroquinolones, aminoglycosides, and ethionamide for Xpert MTB/XDR), our system increased the sensitivity and negative predictive values on the independent test set compared to the expected performance of these assays, at a small cost in specificity and positive predictive value. There are several explanations, including that the assays only look at eight genes and promoter regions, and exclude rare variants therein,^5^ while the machine learning system is able to explore genome-wide features, leverage interactions between features, and assess lineage and genetic background through genome-wide features. While we did not compare our approach to the new targeted next generation sequencing (tNGS) assays such as Deeplex MycTB (Genoscreen),^20^ the ability to harness genome-wide information comprises a theoretical advantage over such assays.

Potential benefits of our approach include the ability to update and train automatically as new resistant samples are added. As resistance to drugs like bedaquiline emerges, this could help avoid the expensive multi-phase multi-year period required for the development of molecular assays,^21,22^ or the need to update catalogues through expert review.^16^ The U.S. Food and Drug Administration (FDA) recently released a regulatory framework for ’live’ modifications to artificial intelligence and machine learning-based software as a medical device,^23^ and has recently provided clearance or approval for several such diagnostic devices,^24^ paving the way for clinical implementation and dissemination. Further advantages of our approach include that predictions are made for all isolates, avoiding the exclusion of 4-10% of samples with unknown mutations in candidate genes.^25^ The use of *k*-mers from sequencing reads allows for genome-wide analysis whilst avoiding some of the vulnerabilities associated with read mapping or variant calling that can affect resistance prediction.^26^ Whereas a *vcf* file filters out sites called with low confidence, our machine learning approach uses all *k*-mers from reads as training features. Finally, by using an interpretable supervised machine learning algorithm, we provide a list of features used for prediction, which in turn can be hypothesis-generating in the search for causal mutations.

We note several limitations to our study. First, we were unable to validate essential agreement of MICs outside of the CRyPTIC dataset, due to the unavailability of such data (CRyPTIC is the first study to systematically collect MIC data across so many drugs). While we validate against binary phenotypes derived from MGIT, a priority of future work should be to perform external validation of MICs when such data becomes available. Second, there were a very small number of isolates phenotypically resistant to bedaquiline, linezolid, delamanid, and clofazimine in the CRyPTIC dataset. Consequently, we report MIC data, and both a train-test approach and cross-validation of models, tested on each site and trained on all other sites, but error bars remain large. A third limitation is the use of a previously published literature-derived catalogue, rather than the more recently published WHO-endorsed catalogue,^16^ because the WHO catalogue was developed using samples from both the CRyPTIC and independent datasets. Fourth, we report the performance of GeneXpert in silico, but clinical performance of the actual method might differ.

In summary, this study demonstrates that WGS can be combined with simple machine learning algorithms to provide MIC predictions for many drugs recommended for the treatment of susceptible and of MDR-TB. This study shows how a composite machine learning system could be used to help guide therapy, whilst being straightforward to updated as increasing numbers of resistant samples to new and repurposed drugs are collected. Further data and external validation are required before clinical implementation.

## Methods

### Study design

We performed a training, validation, and external testing study of a machine learning system to predict minimum inhibitory concentration (MIC) to 13 antituberculosis antibiotics using whole-genome sequencing data (WGS). We trained and tested the system on 10,859 isolates from 11 laboratories in 22 countries collected by the CRyPTIC consortium. Phenotypes were determined using the UKMYC broth microdilution system.^15^ We then assessed how this system, trained on UKMYC-derived phenotypes, would perform against a commonly used DST method in independent samples. For this we made predictions for an external set of isolates used to derive the WHO catalogue of drug-resistant mutations.^15^ We selected only those samples that had been phenotypically characterised by Mycobacteria Growth Indicator Tube (MGIT), namely 15,239 *M. tuberculosis complex* isolates from 22 countries (Table 1, Table S9).

### Genotypic data

#### Whole-genome sequencing and k-mer generation

All isolates were whole-genome sequenced using Illumina next-generation sequencing, with sequencing protocols varying between sites as previously described.^15^ Sequencing reads were trimmed and mapped to the reference genome H37Rv, and variants called using Clockwork (v0.8.3), a bespoke processing pipeline built for CRyPTIC and optimised to detect both single nucleotide polymorphisms and indels. Prior to mapping and calling, raw nucleotide *k*-mers from sequencing reads were set aside for training the machine learning predictor. Each read was decomposed into a series of 31-mers, in line with the standard for bacteria, and for feasibility reasons. While small values of *k* might generate *k*-mers that ambiguously map to many genome loci, large values of *k* will yield very specific *k*-mers that only occur in a few genes. Several studies found a *k*-mer length of 31 to be optimal for bacterial genome assembly^27,28^. 31-mers also offer considerable specificity and a manageable memory footprint^27,28^, as they are the longest *k*-mer length that can be efficiently represented on a 64-bit machine. They have also been demonstrated to be superior to predict AMR with an AdaBoost model. Since *M. tuberculosis* is highly conserved, requiring fewer *k*-mer counts for a given specificity^28^, *k*-mer counts of 31 are most appropriate for this analysis.

#### K-mer frequency correction and pattern generation

*k*-mers were generated in random order, and then reordered by alphabetical order of base pairs to facilitate analysis. While most *k*-mers are present more than 20 times in an isolate, some are present fewer than 5 times (Figure S4). This is likely the result of sequencing errors. One of the disadvantages of using *k*-mers from sequencing reads, as opposed to assembled genomes, is the lack of any error processing. For a given isolate, the frequency distribution of *k*-mer frequencies possesses two peaks: one at a frequency of 1, representing sequencing errors and *k*-mers present once, and another at a frequency of between 100 and 150, representing the true mode frequency. This is illustrated in Figure S4 where the *k*-mer distributions of fifty isolates are displayed, highlighting the variety of mode frequencies and standard deviations, and the presence of a second peak at a frequency of 195 corresponding to sequencing errors. To reduce the influence of these errors on our analysis, all *k*-mers present five times or fewer were removed from the dataset.

#### Pattern features and k-mer matrix generation

Given the number of unique *k*-mers in any dataset is several orders of magnitude greater than the number of samples, further processing is necessary to allow for efficient model training. We combined all groups of *k*-mers that always appear together across all isolates (that is, are present or absent in the same isolates) into a single feature. We refer to features that represent a combination of *k*-mers with the same presence or absence pattern as ’patterns’ of *k*-mers. This procedure is a form of lossless compression, as no data is lost in the process of shrinking the feature space, and was originally described by Earle et al^29^. We used a custom code developed by Earle and Wilson^29^ to perform this pattern compression. First, all *k*-mer files are opened sequentially, and a single list of all unique *k*-mers present in the dataset is generated. Second, *k*-mers are combined in groups, and the pattern of presence or absence for all *k*-mers in the group is generated by interrogating each file successively. The *k*-mers present below a specified frequency threshold of 5 are discarded. Third, groups of k-mers generated in step 2 are combined, and all *k*-mers that follow the exact same pattern of presence or absence across the dataset are combined into a single pattern. The result is a list of patterns, and a key associating each pattern with a list of *k*-mers, allowing for feature analysis. Despite the effective and lossless compression provided by pattern combination, further file compression is required to generate *k*-mer matrices that can be effectively processed by machine learning algorithms. All *k*-mer feature matrices were stored and read using the Hierarchical Data Format (HDF). HDF is a file format system designed and developed to facilitate the management and storage of large data files, with less disk space and faster retrieve.

### Phenotypic data

#### Minimum /nhibitory Concentration phenotypes

Phenotypic drug susceptibility testing (DST) for the CRyPTIC training and test set was performed across all sites using a standard protocol described elsewhere.^15^ Briefly, samples were subcultured and inoculated into 96-well broth microdilution plates containing 13 drugs; the plates were designed by the CRyPTIC consortium and manufactured by Thermo Fisher Inc., U.K. Between 5 and 10 doubling dilutions were used for each drug, and minimum inhibitory concentrations (MIC) for each were read after 14 days using three methods for quality assurance. MICs were converted to predictions of resistance or susceptibility using epidemiological cutoffs (ECOFFs).^15^ As the plate design was modified during the study, the intersect of both plates was used as the MIC phenotype, and concentrations outside both were right censored or left censored as appropriate. Phenotypic DST for the external test set used the BACTEC MGIT 960 system.

#### Phenotype processing

MIC values are both left and right censored. If the bacteria cannot grow in any of the wells, then the MIC must be lower or equal to that of the well with the lowest concentration and the value is left censored. If the bacteria can grow in all the wells, then the MIC must be greater than the highest well concentration and the value is right-censored. Several options can be explored to manage censored data when preparing the label vector. The first option would be to remove all censored phenotypes from the dataset. However, this would lead to a majority of samples being lost (>80% of isolates for rifabutin (RFB)), especially since left censorship is expected in susceptible wild-type isolates when a drug is effective. The second option would be to replace all censored values with the value of the previous or next dilution. Finally, a maximum likelihood estimate could be used by fitting a multi-regression model to the data, preserving the uncertainty of censored MICs.^30^. The challenge of censorship is compounded by the fact that the CRyPTIC dataset includes data from two different phenotypic plates, UKMYC5 and UKMYC6, each censored at different concentrations for each drug. To resolve the discordance between both plates, we only considered the intersection of both - concentrations that were present on both plates - and right or left censored the concentrations present on only one of the two plates (Table S4A). For simplicity and scalability, we then converted the censored values into the value of the previous or next dilution - where >1.6mg/L becomes 3.2mg/L. Finally, to account for the doubling dilutions and the sequential nature of MICs, we converted each value into its binary logarithm log_2_MIC. The resulting vector was used as the regression label vector.

### Model training

#### Hyperparameter tuning

For each model, we used a combination of best practices from the literature, grid search, and random search hyperparameter optimisation. In most cases we would use random search first to find the most impactful hyperparameters and initial values, and grid search for fine-tuning. We set aside an independent set of 1,000 isolates exclusively for hyperparameter tuning to avoid data leakage. We used 5-fold cross-validation with custom mean squared error scoring function. For random search, between 20 and 100 iterations were computed, depending on available computing power and time. In order to correct for class imbalance when evaluating models and performing feature analysis, we use stratified *k*-fold cross-validation for hyperparameter tuning. Samples are divided into *k* folds. One fold is used as the test set, while every other fold is used as the training set. Model performance and feature weights are computed, and the training starts again, using a different fold as the test set. This is repeated *k* times, until each fold has been used as a test set. After this process is complete, each sample has been used in the test set exactly once.

#### M/C prediction

DST for each sample was predicted using a *k*-mer-based, hypothesis-free, genome-wide supervised machine learning algorithm. Raw nucleotide *k*-mers (*k*=31) from sequencing reads (i.e. prior to mapping or assembly) were used as features. A total of 1.9 x 10^9^ individual *k*-mers were considered. Where <5 *k*-mers were identified for an isolate, these were considered sequencing errors (Figure S4). We merged features across patterns,^29^ performed feature selection using the F-test applied to MICs, and trained an optimised tree-based extreme gradient boosting method to allow for rapid training, testing, and feature interpretation. After training, the top features relevant to each prediction were mapped to H37Rv using bowtie2 for detailed feature analysis (Figure S4). Youden’s J statistic was applied to derive the operating threshold of the system.

#### Computation

Given the large dataset size and high number of features, we explored ways to increase the speed of model training and computer memory to allow for training across the entire CRyPTIC dataset. Extreme gradient boosting (XGB) methods generate an ensemble of learners sequentially, and not in parallel, with each new predictor attempting to correct the errors of its predecessors by minimising the residual function. As such, trees cannot be developed in parallel. However, the model’s system design enables it to compute in parallel through the use of a block structure. As the most consuming part of tree learning is getting the data into sorted order, XGB stores the data in in-memory units called blocks. Different blocks may be distributed across cores and machines, or even out-of-core. A full description of the block design is presented in the first XGB implementation paper^31^. All computations for machine learning training on *k*-mers were performed on remote cluster CPU nodes. We used the Biomedical Research Computer (BMRC) cluster of Oxford University, which includes over 7,000 CPU cores and 7 PB (one million GB) of fast, shared storage serving data at up to 30 GB/s. All computers run a Linux operating system. All Python jobs were sent to the cluster via ssh and a custom Univa Grid Engine scheduler. The cluster is composed of a series of computers (or nodes), with each node composed of cores. For processing the entire CRyPTIC dataset, we used a set of 3 Intel Ivybride Node with 48 cores per node and 41.25 GB of memory per core, for a total of 2 TB of memory available. We used the array job function when it was required to send hundreds of jobs across methods, drugs, and sample sizes in parallel.

#### Train-test

Performance on the 25% CRyPTIC test set was estimated by training the system on the 75% CRyPTIC samples not included in it. Performance on the independent test set was generated by training the system on the entire CRyPTIC dataset. P-values were calculated using McNemar chi-square test. We benchmarked the performance of the mutation catalogue and machine learning system against the expected performance of Xpert® MDR/RIF and Xpert® XDR (Cepheid, Sunnyvale, U.S.), based on the targets they probe (Table S13). ‘Indeterminate’ predictions by the catalogue where a novel variation is seen in a candidate gene, were counted as susceptible for the purpose of the analysis. Finally, we simulated negative predictive values for each drug for different prevalence values of resistance. For each drug, we selected 138 samples at random to generate data sets with a percentage prevalence of resistance for every 1% between 1% and 49%, and repeated this 100 times. 138 corresponds to twice the number of isolates resistant to bedaquiline, the drug with the smallest resistance prevalence (69 resistant isolates).

## Supporting information

Tables_all

## Ethics

Approval for CRyPTIC study was obtained by Taiwan Centers for Disease Control IRB No. 106209, University of KwaZulu Natal Biomedical Research Ethics Committee (UKZN BREC) (reference BE022/13), University of Liverpool Central University Research Ethics Committees (reference 2286), Institutional Research Ethics Committee (IREC) of The Foundation for Medical Research, Mumbai (Ref nos. FMR/IEC/TB/01a/2015 and FMR/IEC/TB/01b/2015), Institutional Review Board of P.D. Hinduja Hospital and Medical Research Centre, Mumbai (Ref no. 915-15-CR [MRC]), scientific committee of the Adolfo Lutz Institute (CTC-IAL 47-J / 2017) and the Ethics Committee (CAAE: 81452517.1.0000.0059) and Ethics Committee review, Universidad Peruana Cayetano Heredia (Lima, Peru) and LSHTM (London, UK).

### Members of the CRyPTIC consortium (in alphabetical order)

Correspondence to: Alexander S Lachapelle

Ivan Barilar^29^, Simone Battaglia^1^, Emanuele Borroni^1^, Angela P Brandao^2,3^, Alice Brankin^4^, Andrea Maurizio Cabibbe^1^, Joshua Carter^5^, Daniela Maria Cirillo^1^, Pauline Claxton^6^, David A Clifton^4^, Ted Cohen^7^, Jorge Coronel^8^, Derrick W Crook^4^, Viola Dreyer^29^, Sarah G Earle^4^, Vincent Escuyer^9^, Lucilaine Ferrazoli^3^, Philip W Fowler^4^, George Fu Gao^10^, Jennifer Gardy^11^, Saheer Gharbia^12^, Kelen T Ghisi^3^, Arash Ghodousi^1,13^, Ana Lu za Gibertoni Cruz^4^, Louis Grandjean^33^, Clara Grazian^14^, Ramona Groenheit^44^, Jennifer L Guthrie^15,16^, Wencong He^10^, Harald Hoffmann^17,18^, Sarah J Hoosdally^4^, Martin Hunt^19,4^, Zamin Iqbal^19^, Nazir Ahmed Ismail^20^, Lisa Jarrett^21^, Lavania Joseph^20^, Ruwen Jou^22^, Priti Kambli^23^, Rukhsar Khot^23^, Jeff Knaggs^19,4^, Anastasia Koch^24^, Donna Kohlerschmidt^9^, Samaneh Kouchaki^4,25^, Alexander S Lachapelle^4^, Ajit Lalvani^26^, Simon Grandjean Lapierre^27^, Ian F Laurenson^6^, Brice Letcher^19^, Wan-Hsuan Lin^22^, Chunfa Liu^10^, Dongxin Liu^10^, Kerri M Malone^19^, Ayan Mandal^28^, Mikael Mansjo^44^, Daniela Matias^21^, Graeme Meintjes^24^, Flavia F Mendes^3^, Matthias Merker^29^, Marina Mihalic^18^, James Millard^30^, Paolo Miotto^1^, Nerges Mistry^28^, David AJ Moore^31,8^, Kimberlee A Musser^9^, Dumisani Ngcamu^20^, Nhung N Hoang^32^, Stefan Niemann^29, 48^, Kayzad Soli Nilgiriwala^28^, Camus Nimmo^33^, Nana Okozi^20^, Rosangela S Oliveira^3^, Shaheed Vally Omar^20^, Nicholas I Paton^34^, Timothy EA Peto^4^, Juliana MW Pinhata^3^, Sara Plesnik^18^, Zully M Puyen^35^, Marie Sylvianne Rabodoarivelo^36^, Niaina Rakotosamimanana^36^, Paola MV Rancoita^13^, Priti Rathod^21^, Esther Robinson^21^, Gillian Rodger^4^, Camilla Rodrigues^23^, Timothy C Rodwell^37,38^, Aysha Roohi^4^, David Santos-Lazaro^35^, Sanchi Shah^28^, Thomas Andreas Kohl^29^, E Grace Smith^21,12^, Walter Solano^8^, Andrea Spitaleri^1,13^, Philip Supply^39^, Utkarsha Surve^23^, Sabira Tahseen^40^, Nguyen Thuy Thuong Thuong^32^, Guy Thwaites^32,4^, Katharina Todt^18^, Alberto Trovato^1^, Christian Utpatel^29^, Annelies Van Rie^41^, Srinivasan Vijay^42^, Timothy M Walker^4,32^, A Sarah Walker^4^, Robin M Warren^43^, Jim Werngren^44^, Maria Wijkander^44^, Robert J Wilkinson^45,46,26^, Daniel J Wilson^4^, Penelope Wintringer^19^, Yu-Xin Xiao^22^, Yang Yang^4^, Zhao Yanlin^10^, Shen-Yuan Yao^20^, Baoli Zhu^47^

#### Institutions

1 IRCCS San Raffaele Scientific Institute, Milan, Italy

2 Oswaldo Cruz Foundation, Rio de Janeiro, Brazil

3 Institute Adolfo Lutz, Sao Paulo, Brazil

4 University of Oxford, Oxford, UK

5 Stanford University School of Medicine, Stanford, USA

6 Scottish Mycobacteria Reference Laboratory, Edinburgh, UK

7 Yale School of Public Health, Yale, USA

8 Universidad Peruana Cayetano Heredia, Lima, Peru

9 Wadsworth Center, New York State Department of Health, Albany, USA

10 Chinese Center for Disease Control and Prevention, Beijing, China

11 Bill & Melinda Gates Foundation, Seattle, USA

12 UK Health Security Agency, London, UK

13 Vita-Salute San Raffaele University, Milan, Italy

14 University of Sydney, Australia

15 The University of British Columbia, Vancouver, Canada

16 Public Health Ontario, Toronto, Canada

17 SYNLAB Gauting, Munich, Germany

18 Institute of Microbiology and Laboratory Medicine, IMLred, WHO-SRL Gauting, Germany

19 EMBL-EBI, Hinxton, UK

20 National Institute for Communicable Diseases, Johannesburg, South Africa

21 Public Health England, Birmingham, UK

22 Taiwan Centers for Disease Control, Taipei, Taiwan

23 Hinduja Hospital, Mumbai, India

24 University of Cape Town, Cape Town, South Africa

25 University of Surrey, Guildford, UK

26 Imperial College, London, UK

27 Universite de Montreal, Canada

28 The Foundation for Medical Research, Mumbai, India

29 Research Center Borstel, Borstel, Germany

30 Africa Health Research Institute, Durban, South Africa

31 London School of Hygiene and Tropical Medicine, London, UK

32 Oxford University Clinical Research Unit, Ho Chi Minh City, Viet Nam

33 University College London, London, UK

34 National University of Singapore, Singapore

35 Instituto Nacional de Salud, Lima, Peru

36 Institut Pasteur de Madagascar, Antananarivo, Madagascar

37 FIND, Geneva, Switzerland

38 University of California, San Diego, USA

39 Univ. Lille, CNRS, Inserm, CHU Lille, Institut Pasteur de Lille, U1019 - UMR 9017 - CIIL - Center for Infection and Immunity of Lille, F-59000 Lille, France

40 National TB Reference Laboratory, National TB Control Program, Islamabad, Pakistan

41 University of Antwerp, Antwerp, Belgium

42 University of Edinburgh, Edinburgh, UK

43 SAMRC Centre for Tuberculosis Research, Stellenbosch University, Cape Town, South Africa

44 Public Health Agency of Sweden, Solna, Sweden

45 Wellcome Centre for Infectious Diseases Research in Africa, Cape Town, South Africa

46 Francis Crick Institute, London, UK

47 Institute of Microbiology, Chinese Academy of Sciences, Beijing, China

48 German Center for Infection Research (DZIF), Hamburg-Lubeck-Borstel-Riems, Germany

### Additional authors contributing to the CRyPTIC consortium (in alphabetical order)

Irena Arandjelovic^1^, Anna Barbova^2^, Maxine Caws^3^, If\aki Comas^4^, Roland Diel^5^, Carla Duncan^6^, Sarah Dunstan^7^, Maha Farhat^8^, Margaret M Fitzgibbon^9^, Victoria Furi6^10^, Jennifer Gardy^11^, Jennifer Guthrie^6^, Dang Thi Minh Ha^12^, Kathryn Holt^13^, Michael Inouye^14^, Frances B Jamieson^6^, SM Mostofa Kamal^15^, Julianne V Kus^6^, Vanessa Mathys^16^, Rick Twee-Hee Ong^17^, Youwen Qin^7,14^, Thomas R Rogers^9,19^, Gian Maria Rossolini^20^, Emma Roycroft^9^, Vitali Sintchenko^21^, Alena Skrahina^22^, Yik Ying Teo^17^, Phan Vuong Khac Thai^12^, Dick van Soolingen^23^, Mark Wilcox^24^, Matteo Zignol^25^

#### Institutions

1 University of Belgrade, Belgrade, Serbia

2 National Institute of phthisiology and pulmonology NAMS Ukraine, Kyiv

3 Liverpool School of Tropical Medicine, United Kingdom

4 Biomedicine Institute of Valencia IBV-CSIC, Spain

5 University Medical Hospital Schleswig-Holstein, Germany

6 Public Health Ontario, Toronto, Canada

7 University of Melbourne, Australia

8 Harvard Medical School, Boston, USA

9 Irish Mycobacteria Reference Laboratory, Dublin, Ireland

10 Universitat de Valencia, Spain

11 Bill & Melinda Gates Foundation, Seattle, USA

12 Pham Ngoc Thach Hospital, Ho Chi Minh City, Vietnam

13 Monash University, Melbourne, Australia

14 Baker Institute, Melbourne, Australia

15 National Institute of Diseases of the Chest and Hospital, Dhaka, Bangladesh

16 Sciensano, Belgian reference laboratory for M. tuberculosis

17 National University of Singapore, Singapore

19 Trinity College Dublin, Ireland

20 Careggi University Hospital, Florence, Italy

21 University of Sydney, Australia

22 Republican Scientific and Practical Centre for Pulmonology and TB, Minsk, Belarus

23 National Institute for Public Health and the Environment, Bilthoven, The Netherlands

24 Leeds Teaching Hospital NHS Trust, Leeds, United Kingdom

25 World Health Organization, Geneva

## Author contributions

DAC, DMC, DWC, HH, SJH, NAI, NM, DM, SN, TEAP, CR, GS, PS, GT, ASW, TMW, DJW, ZY contributed to high-level CRyPTIC study design. ASL, TMW, DWC, TEAP, ASW, PWF, DAC designed the specifics of this study. MH, JK, ZI and PWF retrieved and processed genotypic data including k-mers. PWF retrieved and processed phenotypic data. ASL, DAC, TMW, SK, YY, PWF, developed the machine learning system. All other authors contributed to the generation of data. ASL and TMW performed all the analysis. ASL and TMW wrote the manuscript with all authors offering feedback.

## Acknowledgements & funding

This work was supported by Wellcome Trust/Newton Fund-MRC Collaborative Award (200205/Z/15/Z); and Bill & Melinda Gates Foundation Trust (OPP1133541). Oxford CRyPTIC consortium members are funded/supported by the National Institute for Health Research (NIHR) Oxford Biomedical Research Centre (BRC), and the National Institute for Health Research (NIHR) Health Protection Research Unit in Healthcare Associated Infections and Antimicrobial Resistance, a partnership between Public Health England and the University of Oxford. The views expressed are those of the authors and not necessarily those of the NHS, the NIHR, Public Health England or the Department of Health and Social Care. J.M. is supported by the Wellcome Trust (203919/Z/16/Z). Z.Y. is supported by the National Science and Technology Major Project, China Grant No. 2018ZX10103001. K.M.M. is supported by EMBL’s EIPOD3 programme funded by the European Union’s Horizon 2020 research and innovation programme under Marie Skfodowska Curie Actions. T.C.R. is funded in part by funding from Unitaid Grant No. 2019-32-FIND MDR. R.S.O. is supported by FAPESP Grant No. 17/16082-7. L.F. received financial support from FAPESP Grant No. 2012/51756-5. B.Z. is supported by the National Natural Science Foundation of China (81991534) and the Beijing Municipal Science & Technology Commission (Z201100005520041). N.T.T.T. is supported by the Wellcome Trust International Intermediate Fellowship (206724/Z/17/Z). G.T. is funded by the Wellcome Trust. R.W. is supported by the South African Medical Research Council. J.C. is supported by the Rhodes Trust and Stanford Medical Scientist Training Program (T32 GM007365). T.C. has received grant funding and salary support from US NIH, CDC, USAID and Bill and Melinda Gates Foundation. L.G. was supported by the Wellcome Trust (201470/Z/16/Z), the National Institute of Allergy and Infectious Diseases of the National Institutes of Health under award number 1R01AI146338, the GOSH Charity (VC0921) and the GOSH/ICH Biomedical Research Centre (www.nihr.ac.uk). A.L. is supported by the National Institute for Health Research (NIHR) Health Protection Research Unit in Respiratory Infections at Imperial College London. S.G.L. is supported by the Fonds de Recherche en Sante du Quebec. C.N. is funded by Wellcome Trust Grant No. 203583/Z/16/Z. A.V.R. is supported by Research Foundation Flanders (FWO) under Grant No. G0F8316N (FWO Odysseus). G.M. was supported by the Wellcome Trust (098316, 214321/Z/18/Z, and 203135/Z/16/Z), and the South African Research Chairs Initiative of the Department of Science and Technology and National Research Foundation (NRF) of South Africa (Grant No. 64787). The funders had no role in the study design, data collection, data analysis, data interpretation, or writing of this report. The opinions, findings and conclusions expressed in this manuscript reflect those of the authors alone. A.B. is funded by the NDM Prize Studentship from the Oxford Medical Research Council Doctoral Training Partnership and the Nuffield Department of Clinical Medicine. D.J.W. is supported by a Sir Henry Dale Fellowship jointly funded by the Wellcome Trust and the Royal Society (Grant No. 101237/Z/13/B) and by the Robertson Foundation. A.S.W. is an NIHR Senior Investigator. T.M.W. is a Wellcome Trust Clinical Career Development Fellow (214560/Z/18/Z). A.S.L. is supported by the Rhodes Trust. R.J.W. receives funding from the Francis Crick Institute which is supported by Wellcome Trust, (FC0010218), UKRI (FC0010218), and CRUK (FC0010218). The computational aspects of this research were supported by the Wellcome Trust Core Award Grant Number 203141/Z/16/Z and the NIHR Oxford BRC. Parts of the work were funded by the German Center of Infection Research (DZIF). The Scottish Mycobacteria Reference Laboratory is funded through National Services Scotland. The Wadsworth Center contributions were supported in part by Cooperative Agreement No. U60OE000103 funded by the Centers for Disease Control and Prevention through the Association of Public Health Laboratories and NIH/NIAID grant AI-117312. Additional support for sequencing and analysis was contributed by the Wadsworth Center Applied Genomic Technologies Core Facility and the Wadsworth Center Bioinformatics Core. SYNLAB Holding Germany GmbH for its direct and indirect support of research activities in the Institute of Microbiology and Laboratory Medicine Gauting. N.R. thanks to the Programme National de Lutte contre la Tuberculose de Madagascar.

## Acknowledgements - people

We thank Faisal Masood Khanzada and Alamdar Hussain Rizvi (NTRL, Islamabad, Pakistan), Angela Starks and James Posey (Centers for Disease Control and Prevention, Atlanta, USA), and Juan Carlos Toro and Solomon Ghebremichael (Public Health Agency of Sweden, Solna, Sweden).

## Data and code availability

All data used in this manuscript are publicly available on the European Bioinformatics Institute (http://ftp.ebi.ac.uk/pub/databases/cryptic/).

## Wellcome Trust Open Access

This research was funded in part, by the Wellcome Trust/Newton Fund-MRC Collaborative Award [200205/Z/15/Z]. For the purpose of Open Access, the author has applied a CC BY public copyright licence to any Author Accepted Manuscript version arising from this submission.

This research was funded, in part, by the Wellcome Trust [214321/Z/18/Z, and 203135/Z/16/Z]. For the purpose of open access, the author has applied a CC BY public copyright licence to any Author Accepted Manuscript version arising from this submission.

## Figures and Tables

All primary and supplementary figures and tables are presented in separate files (*Figures.pdf* and *Tables.xlsx*).

---

**Figure S1:**
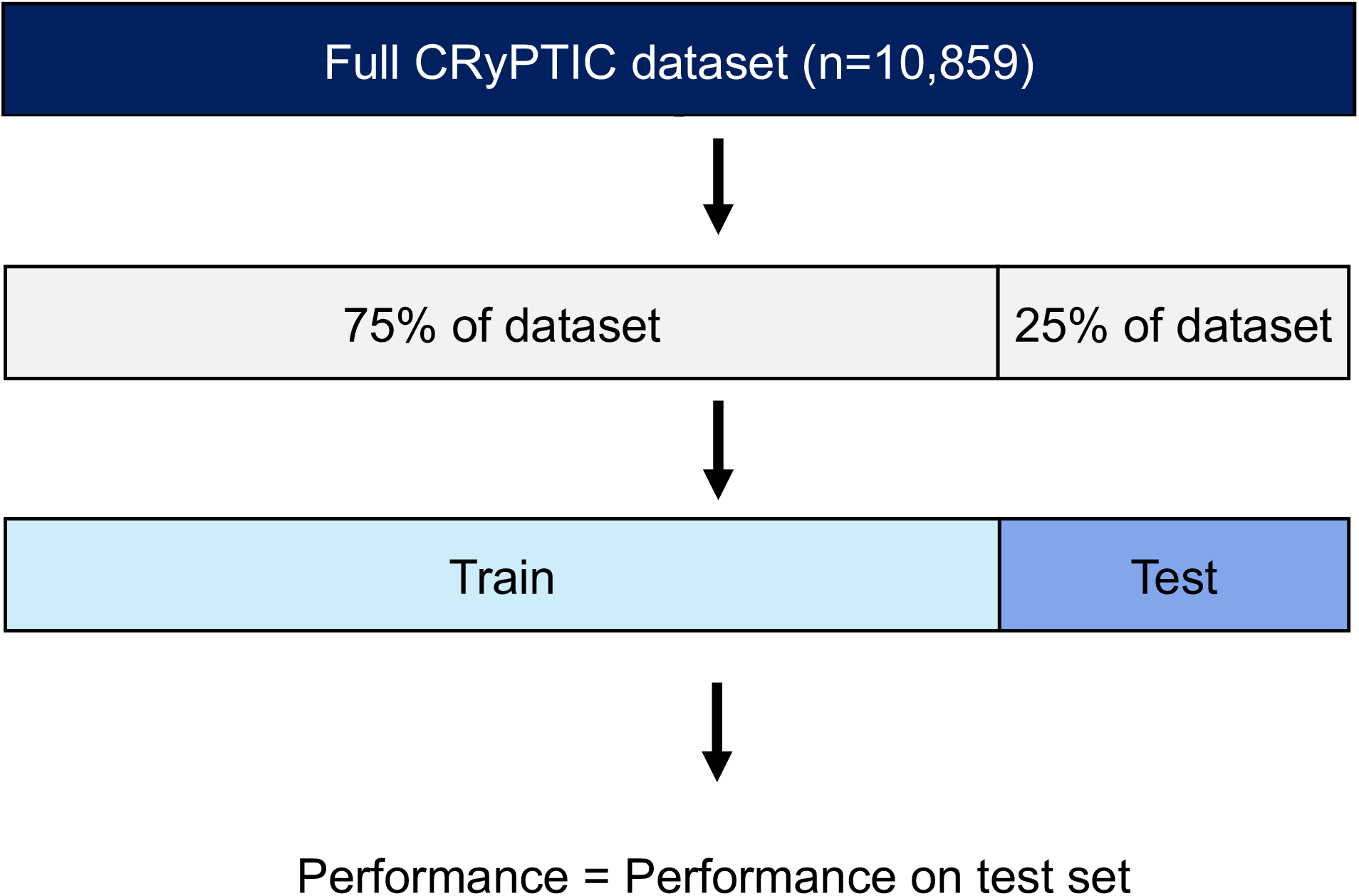
Illustration of training and testing process in CRyPTIC dataset

**Figure S2:**
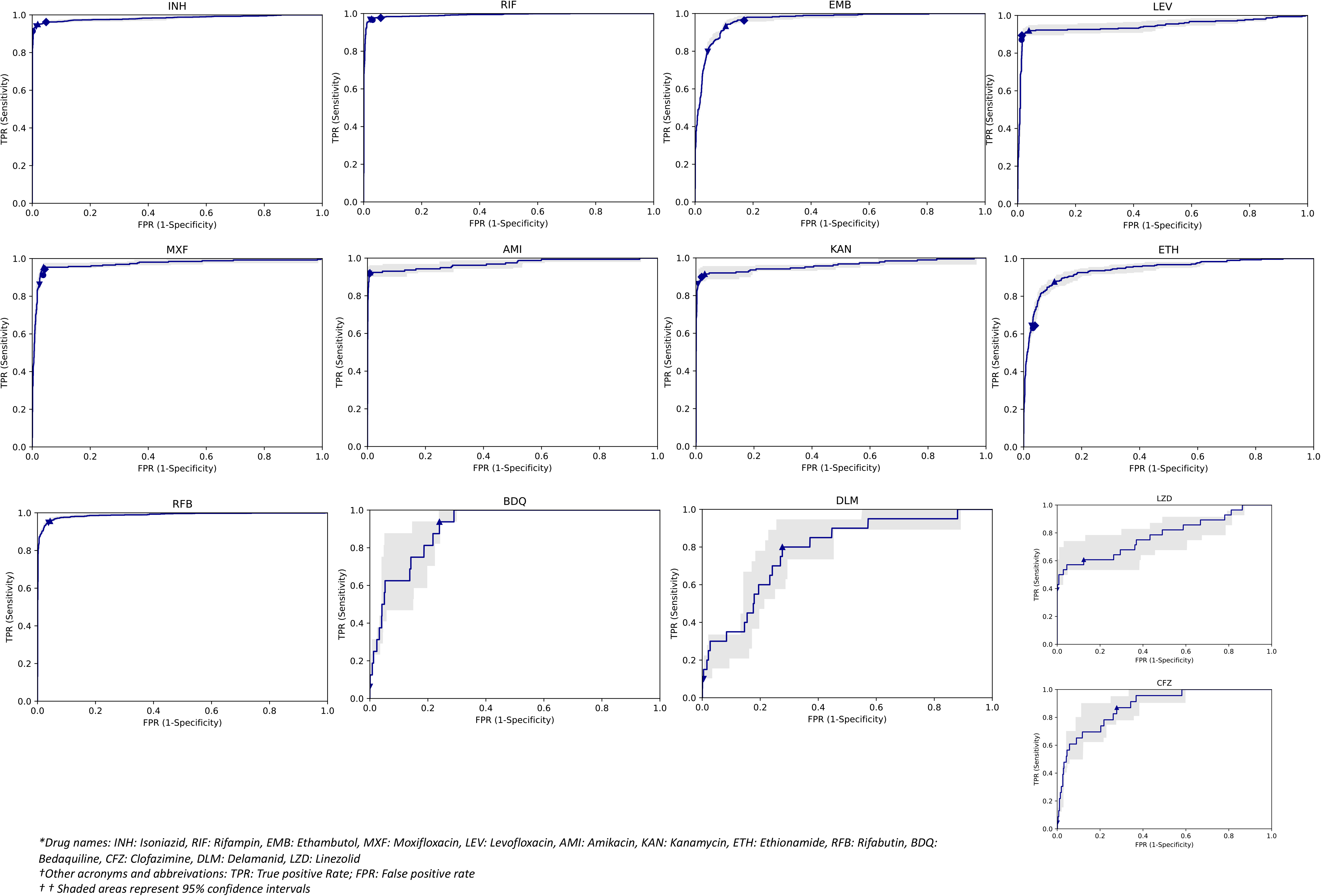
Receiver operating characteristic curves for binary predictions in CRyPTIC dataset

**Figure S3:**
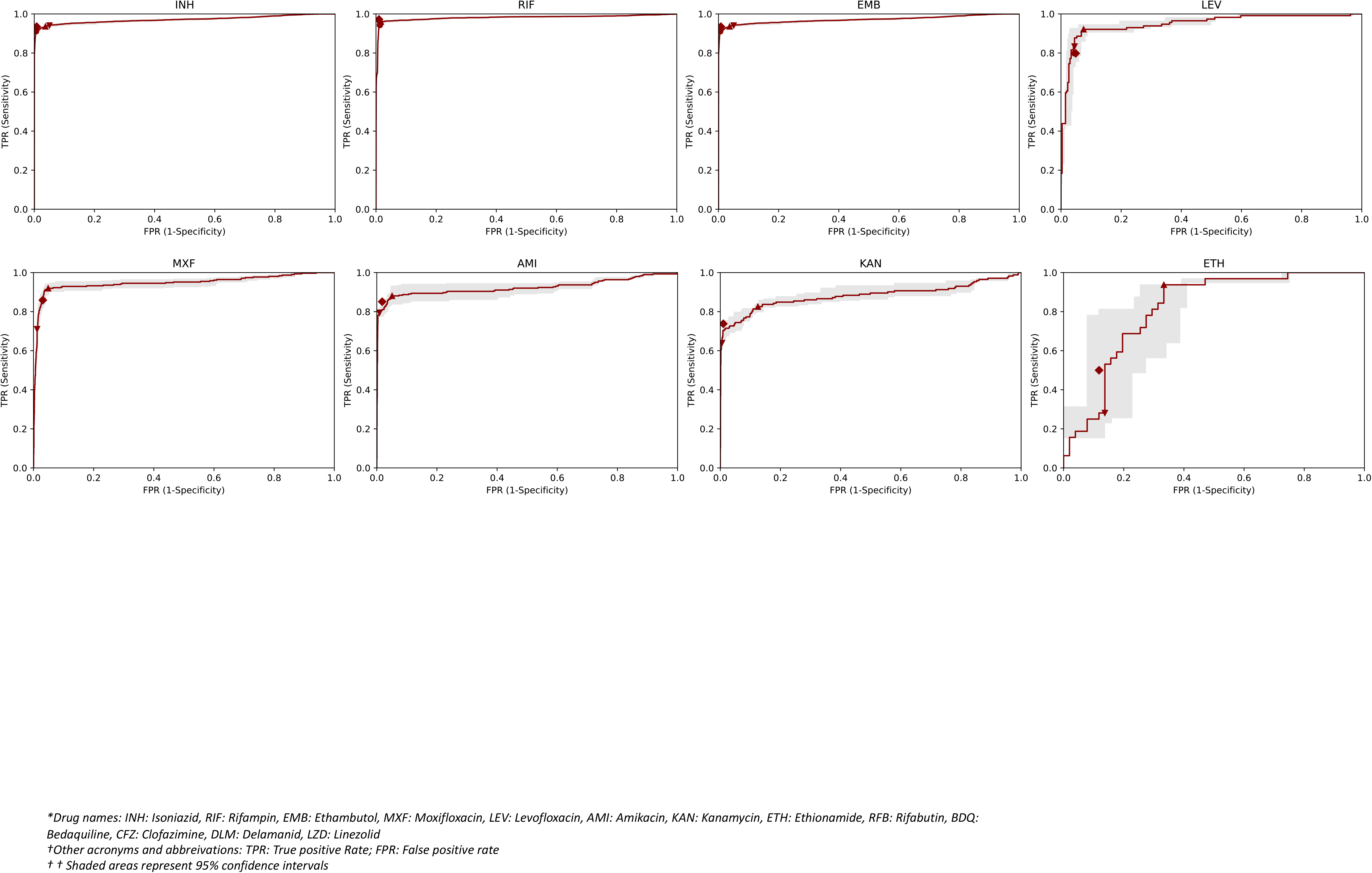
Receiver operating characteristic curves for binary predictions in independent dataset

**Figure S4:**
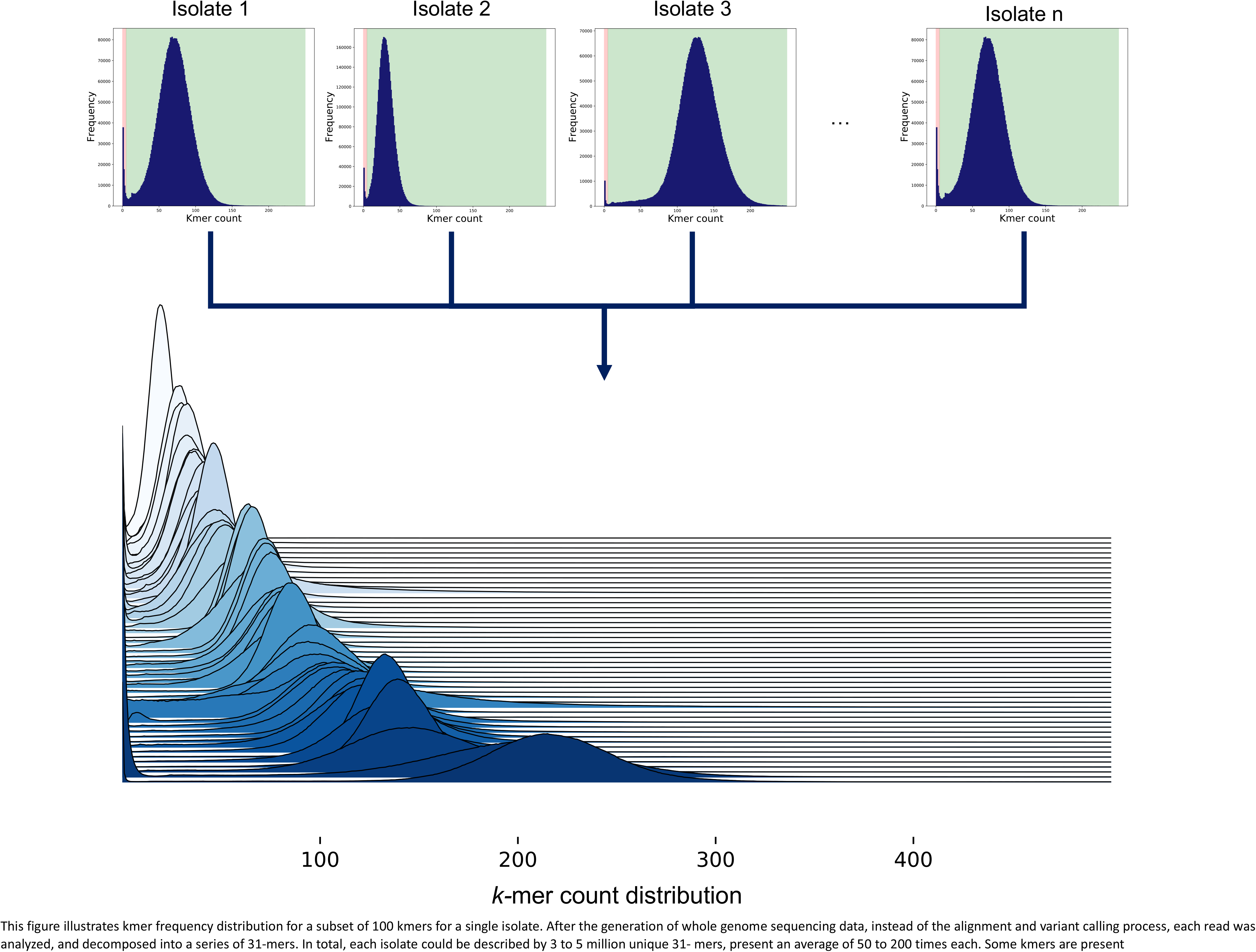
Illustration of kmer distributions across isolates used as features for the machine learning model

**Figure S5:**
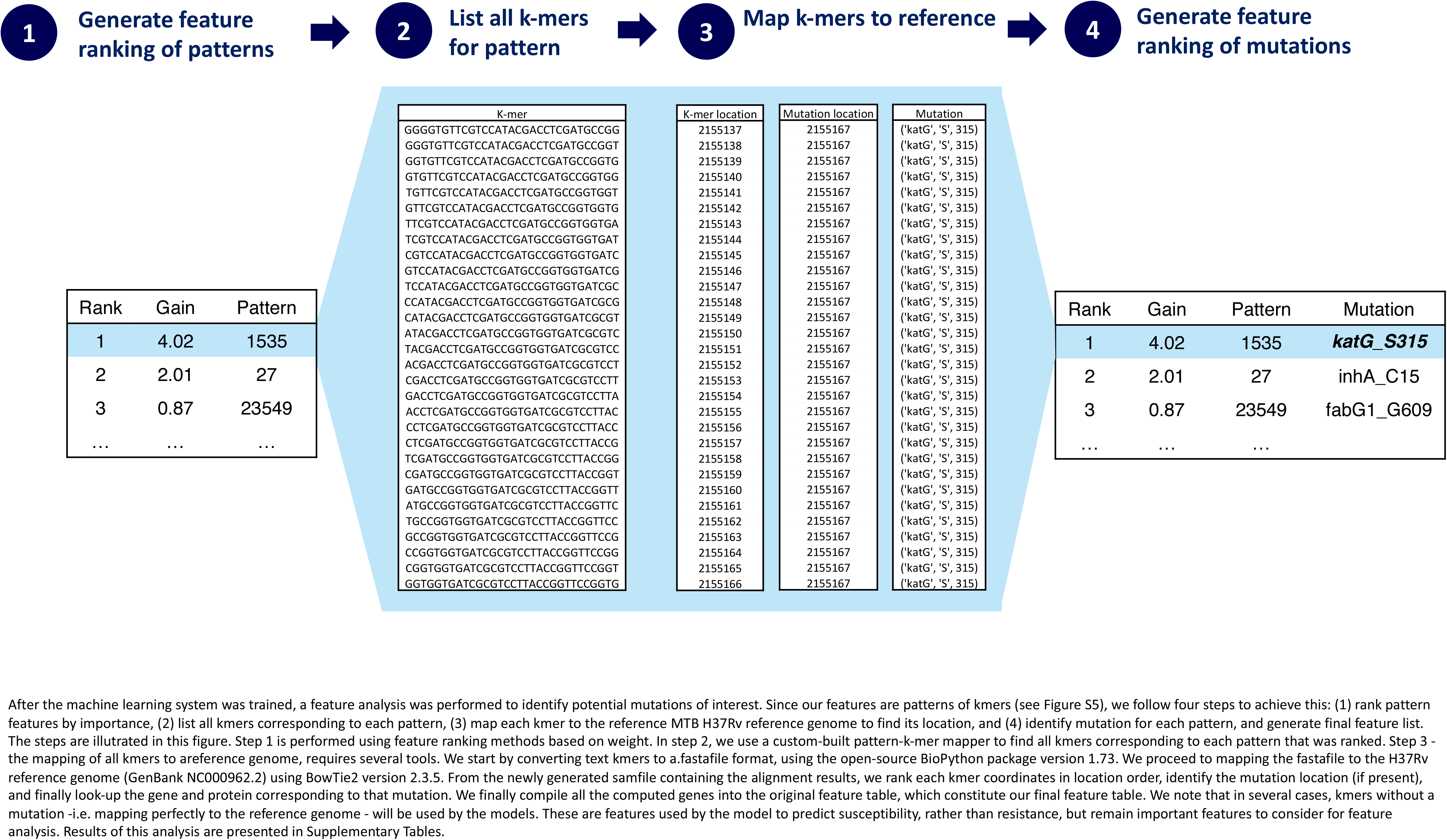
Illustration of the pipeline for kmer feature analysis after machine learning system training

